# Systemic drivers and molecular mechanisms of sarcopenia in aetiology-specific end-stage liver disease

**DOI:** 10.1101/2025.08.21.671433

**Authors:** Thomas Nicholson, Sophie L. Allen, Jonathan I. Quinlan, Amritpal Dhaliwal, Michael Macleod, Joshua Price, Jon Hazeldine, Michael S. Sagmeister, Caitlin Ditchfield, Kirsty C. McGee, Felicity R. Williams, Ahmed M. Elsharkawy, Matthew J. Armstrong, Carolyn A. Greig, Janet M. Lord, Leigh Breen, Simon W. Jones

## Abstract

**Introduction:** Patients with end-stage liver disease (ESLD) often present with sarcopenia, defined as loss of skeletal muscle mass and quality, which is associated with reduced quality of life and increased mortality. However, the molecular mechanisms driving sarcopenia in ESLD are not fully understood and there are currently no therapeutic interventions. This study aimed to identify potential circulating factors contributing to sarcopenia progression in ESLD by assessing their role in driving transcriptomic alterations in skeletal muscle.

**Methods:** Quadriceps muscle tissue, plasma and serum were obtained from ESLD patients (n=24) and age/sex-matched healthy controls (HC; n=18) (clinical trial ID: NCT04734496, ethical approval 18/WM/0167). Total RNA from snap-frozen vastus lateralis muscle biopsies underwent RNA sequencing (Illumina). Serum concentrations of 60 cytokines were profiled by Luminex and ELISA, with comparisons made both between ESLD and HC, and across ESLD aetiologies (alcohol-related, NAFLD, viral hepatitis, other). In vitro, primary human myotubes (from non-ESLD aged donors, NRES #16/SS/0172) were treated with 10% ESLD or HC plasma (24 h, n=6 per group) followed by RNA sequencing (BGI Genomics). Differentially expressed genes (p<0.05, fold-change >1.5) were identified via Qlucore and DESeq2, and pathway analysis performed using Ingenuity (Qiagen). The impact of physiological concentrations of candidate cytokines (IL-1α, GDF-15, HGF) on myotube thickness, differentiation and mitochondrial function was assessed by immunofluorescence microscopy, RT-qPCR and metabolic flux assays.

**Results:** In ESLD muscle, 387 and 225 genes were significantly up- and downregulated compared to HC respectively, with cellular senescence identified as a top dysregulated function. Upstream regulator analysis predicted activation of hepatocyte growth factor (HGF) and interleukin-1 signalling. Subgroup analysis revealed distinct transcriptomic profiles based on disease aetiology. Serum profiling identified 15 cytokines significantly elevated (p<0.05) and 5 reduced (p<0.05) in ESLD, including increased HGF and reduced interleukin-1 receptor antagonist. Stratified analysis also revealed aetiology specific cytokine profiles, with only GDF-15 significantly (P<0.0001) elevated in all groups. 24h ESLD plasma treatment induced 423 differentially expressed genes in human myotubes, which were again associated with significant activation of senescence pathways, with IL-1 identified as a key upstream driver. In vitro, IL-1α, GDF-15, and HGF significantly reduced myotube thickness, nuclear fusion index and perturbed metabolism (Increased glycolysis, impaired oxidative phosphorylation).

**Conclusions:** Collectively, these findings suggest that sarcopenia in ESLD is driven by aetiology-specific mechanisms, highlighting the potential for targeted therapies to improve muscle mass and function.

## Introduction

End stage liver disease (ESLD) is a major healthcare problem, with year on year increases in deaths over the last 15 years (1). One in three patients with ESLD develop sarcopenia, defined as a significant reduction in skeletal muscle mass and/or muscle function (2). Sarcopenia significantly impairs functional ability, impacting quality of life and is associated with an increased risk of falls, hospitalisation and clinical complications such as encephalopathy and infection (3–5). Consequently, ESLD patients with sarcopenia have worse health outcomes post liver transplantation and a 2-fold increase in mortality (6).

Resulting in the decline in muscle mass, sarcopenia reflects a chronic negative imbalance between muscle protein synthesis and protein degradation. In the context of liver disease, current evidence suggests that a chronic state of malnutrition, hyperammonaemia (7), dysregulated muscle protein turnover (8, 9) and reduced amino acid availability (10, 11) may all play a role in the development of sarcopenia (12). However, due to the difficulty in obtaining skeletal muscle tissue from such a clinically vulnerable cohort, there is a scarcity of data regarding the molecular mechanisms that underpin skeletal muscle dysfunction in ESLD. Additionally, whether there are key aetiology-specific mechanisms that drive sarcopenia in ESLD patients has not been described. ESLD is an umbrella term that encompasses different disease aetiologies, typically; Alcohol related liver disease (ArLD), metabolic dysfunction-associated steatotic liver disease (MASLD) and immune associated liver disease (IALD), (encompassing Primary Biliary Cirrhosis, Primary Sclerosing Cholangitis and Autoimmune Hepatitis.). Although sarcopenia is prevalent within all of these disease subgroups (13), they are clinically very distinct populations with considerable differences in both physical characteristics and the underlying pathology. As a result, there is currently poor consensus on the best therapeutic strategy to combat sarcopenia in ESLD patients, especially in an aetiology specific manner. Furthermore, there are no approved pharmacological interventions available that can improve or maintain skeletal muscle mass. Thus, identification of aetiology specific mechanisms of sarcopenia offers a significant opportunity to better inform therapeutic strategies and to develop targeted pharmacological therapeutics to improve skeletal muscle mass and function in these patients.

Here, we have performed the first transcriptomic analysis of skeletal muscle tissue obtained from a deeply phenotyped cohort of ESLD patients, comprised of ArLD, MASLD and IALD aetiologies, to determine universal and aetiology specific mechanisms of skeletal muscle dysfunction in ESLD. Additionally, we have comprehensively profiled circulating cytokines in ESLD patient serum and subsequently performed *ex vivo* experiments with primary human myotubes to examine the role of candidate cytokines in driving sarcopenia across different ESLD aetiologies.

## Methods

### Participants and ethical approval

ESLD patients enrolled in the present study were recruited as part of a larger prospective observational study, namely, the Evaluation of Sarcopenia in Inflammatory Disease (clinical trial ID: NCT04734496), in which the inclusion and exclusion criteria have previously been reported. All ESLD patients had end-stage liver disease (Child_Pugh B/C). Age- and sex-matched control participants had normal liver test results (including blood, risk factor, and imaging data), had no medical comorbidities, and were not taking any medication. Ethical approval was obtained through the Health Research Authority—West Midlands Solihull Research Ethics Committee Authority (REC reference: 18/WM/0167) for the recruitment of patients with ESLD and the local Ethics Committee at the University of Birmingham (ERN_19-0831) for the recruitment of healthy controls. Additionally, for *in vitro* validation experiments, muscle tissue was obtained from patients without ESLD, from patients undergoing total hip arthroplasty (NRES #16/SS/0172). All studies were conducted in accordance with the Declaration of Helsinki, and all participants provided written informed consent.

### Study design

All participants reported to the laboratory in a fasted state (from 06:00 on the day of visit) and were asked to refrain from the consumption of caffeine on the morning of the trial. In addition, participants were asked to refrain from strenuous exercise for 24_h prior to their laboratory visit. Upon arrival, a fasted venous blood sample was obtained for serum/plasma collection. Participants underwent assessments of basic body composition, skeletal muscle mass, and functional strength as previously described () . A skeletal muscle biopsy was obtained from the vastus lateralis of the dominant leg using a Bergström needle 24 and immediately snap frozen in liquid nitrogen. The samples were stored at −80_°C until analysed. Quadricep muscle performance tests were performed post-biopsy to ensure that the biopsy samples were obtained at rest (14). Measures of quadricep muscle mass and intramuscular adipose tissue (IMAT) accumulation were determined from MR images as described previously (14).

### Blood sample processing

To obtain serum, following a 30 min incubation at room temperature, blood samples were centrifuged at 1620 × g for 10_min at 4°C. The upper serum layer was collected and stored at −80_°C until thawing on ice for analysis. To obtain plasma, heparin anti-coagulated whole blood samples were centrifuged at 438 x g at 4°C for 8 min. The upper serum layer was collected and stored at −80_°C until thawing on ice for analysis. Freeze_thaw cycles were avoided.

### Quantification of serum cytokine concentrations

The concentrations of serum cytokines were measured using commercially available Luminex kits (48 plex, #12007283, Bio-Rad, Hertfordshire, UK), (Diabetes 10-Plex #171A7001M, Bio-Rad, Hertfordshire, UK) or individual ELISAs (DY957, DY788; R&D Systems, MN, USA) following manufacturer protocols. All the samples were measured in duplicate.

### Stimulation of human primary myotubes with recombinant proteins and patient plasma

Primary human myoblasts or myotubes (isolation and differentiation methodology provided in Supporting Information) were stimulated with recombinant HGF (230-30011 Cambridge Bioscience, UK), IL-1α (200-LA Biotechne, UK), GDF-15 (8146-GD, Biothechne UK) or patient plasma as detailed in figure legends. To prevent coagulation, 2.11 units of heparin was added to plasma samples, vortexed and incubated for 1 h on ice. Samples were then centrifuged at 1620 × g for 10 min, 4°C and the supernatant collected.

### Quantification of myotube thickness and nuclear fusion index

Quantification of myotube thickness and nuclear fusion index was performed on myotubes cultured in 24-well plates as described in Supporting Information. For quantification of myotube thickness (MTT), 10 images per well were obtained using a x63 objective. For assessment of nuclear fusion index (NFI), 5 images per well were obtained using a x20 objective. Image analysis was carried out using ImageJ software. The MTT of each myotube was calculated by taking the average of 5 measurements obtained along its length. The NFI was defined as the number of nuclei clearly incorporated into myotubes (myotube defined as a desmin-positive structure with 3 or more nuclei) expressed as a proportion of the total visible nuclei in each field of view, and was calculated by processing images with MyoCount (15).

### RNA isolation

Muscle tissue biopsies (∼10_mg) were first homogenized in RLT buffer (Qiagen, Manchester, UK) supplemented with Beta-mercaptoethanol (β-ME) utilizing a Qiagen Tissue Ruptor (Qiagen, Manchester, UK). RNA was then isolated and treated with DNase using a commercially available kit following the manufacturer’s protocol (RNeasy Fibrous Tissue Mini Kit, Qiagen Manchester, UK). For extraction of RNA from primary human myotubes, culture media was removed and cells directly lysed in RLT buffer (Qiagen, Manchester, UK) supplemented with β-ME. RNA was then isolated and treated with DNase using a commercially available kit following the manufacturer’s protocol (RNeasy blood and tissue kit, Qiagen Manchester, UK). The quantity and quality of RNA were measured utilizing a Bioanalyzer (Agilent, CA, USA).

### Bulk RNA sequencing

For muscle tissue samples, library preparation was performed by the Genomics Facility at the University of Birmingham using a QuantSeq 3′ kit (Lexogen), with libraries sequenced on an Illumina NextSeq 500 platform. For primary human myotubes, library preparation and RNA sequencing was performed by BGI genomics. Sequencing read quality checks were performed using fastQC, and reads were trimmed using Trimmomatic. Reads were mapped to the hg38 reference human genome using Star Aligner. Differential expression analysis was determined using Qlucore Omics Explorer and the DESeq2 R Bioconductor package. Pathway analysis was performed utilising Ingenuity pathway analysis (IPA).

### RT-qPCR

mRNA levels of MAFbx, MuRF-1, MYF5, IL-6, MYOD, MYOG and FOXO were determined, relative to the housekeeping gene 18S, using the iTaq™ Universal SYBR® Green One-step kit mastermix (Bio-Rad) and gene-specific primers (Table S2). All reactions had a total volume of 5 µl, containing 5 ng of RNA, and were performed in triplicate. For each RT-qPCR performed, a non-template control comprising of only iTaq™ Universal SYBR® Green One-step mastermix and gene-specific primers was included to ensure no contamination of PCR reagents. Data was acquired using a Bio-Rad sfx cycler (Bio-Rad).

### Metabolic flux analysis

Myotube metabolic flux was measured using the Seahorse analyser XF96 (Agilent). Primary human myoblasts were seeded (2X10^4^) onto assay plates pre-coated with 0.2% gelatin. Once confluent, myoblasts were differentiated for 8 days as described above and then treated with or without recombinant proteins for 24 hours. Full methodology for the metabolic flux assay is provided in Supporting Information.

### Measurement of Myotube Protein synthesis and degradation

Protein synthesis and degradation rates in primary human myotubes were measured *in vitro* replicating a previously described protocol(16). Full methodology for these assays is presented in Supporting Information

### Neuronal culture

Neuronal differentiation was conducted using SH-SY5Y human neuroblastoma cells as previously described (17). Full methodology is provided in Supporting Information.

### Statistical analysis

Data analysis was performed using GraphPad Prism v9. The normality of data was established by performing Shapiro–Wilk analysis. For normally distributed data, statistical significance was determined by performing unpaired t tests or ANOVA with Dunnett’s post hoc tests for datasets with multiple groups. The statistical significance of nonparametric data was assessed by performing a Mann_Whitney U or Kruskal_Wallis test, followed by Dunn’s multiple comparison tests. Spearman’s correlations were performed to test for associations between functional measures and serum cytokine concentrations. A P-value of <0.05 was considered to indicate statistical significance.

## Results

### Patients with ESLD exhibit a sarcopenic phenotype

In comparison to healthy controls, patients with ESLD displayed significantly lower quadricep peak anatomical cross-sectional area (ACSA) (P=0.02, Table S1) and non-dominant peak knee extensor torque (P=0.001). The ESLD group also displayed increased adiposity, with a significantly greater dry body mass index (BMI, corrected for ascites and/or peripheral oedema; P=0.01), dry body weight (P=0.01), waist-to-hip ratio (P<0.0001), body fat mass (P=0.04) and quadricep intramuscular adipose tissue (P_<_0.0001). This was despite no significant difference in age between the healthy controls (49.7_±_15 years, *n*_=_18) and ESLD patients (54.2_±10.7 years, *n*_=_40); (P=0.2). Further characteristics, including body composition, skeletal muscle mass, physical function, blood markers, medications and the clinical profile of ESLD patients are presented in Table S1.

### Patients with ESLD display a modified skeletal muscle transcriptome associated with impaired skeletal muscle growth and metabolic dysfunction compared to healthy individuals

To determine whether patients with ESLD exhibit an altered skeletal muscle transcriptome that may contribute to reduced muscle mass and impaired muscle function (Table S1), we performed bulk RNA sequencing on skeletal muscle tissue biopsies obtained from patients with ESLD (*n=*24) and age and sex matched healthy controls (*n*=18). In total, upon comparing ESLD and control muscle, 612 differentially expressed genes (DEGs) were identified (P<0.05, fold change>1.5) (Figure 1A). The most significant DEGs included upregulation of inhibitors of muscle hypertrophy (e.g DEPTOR) and downregulation of IGFN, a gene recently reported to be associated with myogenesis and musculoskeletal ageing (18, 19) (Figure 1A). Interrogation of canonical signalling pathways using ingenuity pathway analysis (IPA) indicated that DEGs identified in ESLD muscle were associated with significantly reduced oxidative phosphorylation and mTOR signalling, in addition to mitochondrial dysfunction and protein ubiquitination (Figure 1A). Additionally, IPA indicated that senescence of cells and ubiquitination of proteins were amongst the most significantly activated cellular functions (Figure 1C). In contrast, processes crucial to skeletal muscle growth and maintenance, (synthesis of protein, proliferation of muscle cells, translation of protein and contractility of muscle) were significantly suppressed (Figure 1D). Upstream molecules predicted to regulate the differential gene expression observed in the ESLD muscle tissue included activation of negative regulators of cellular growth and proliferation (TP63, KMT2D), protein degradation (SYVN1) and inflammation (PPARA). In contrast, regulators of neural innervation (MYRF) and protein synthesis (EIF4E) were predicted to be inhibited (Figure 1E).

**Figure 1.**
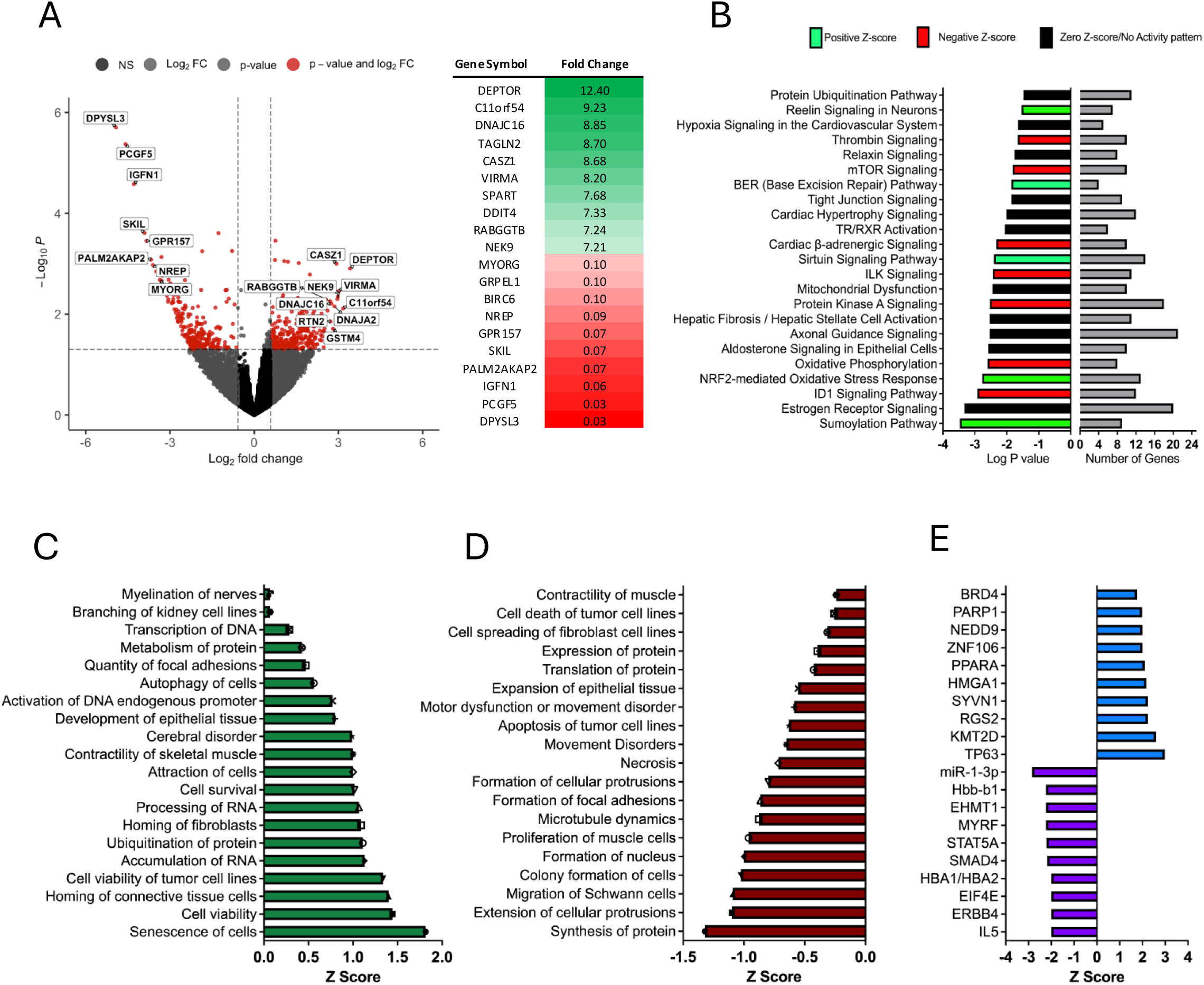
Skeletal muscle tissue from patients with ESLD exhibits an altered transcriptome associated with impaired function. **A.** Volcano plot of DEGs (P<0.05, Fold change >1.5) within the skeletal muscle obtained from ESLD patients vs age- and sex-matched healthy controls. The top 10 DEGs based on fold change are listed to the right. **B.** Top canonical signalling pathways identified in ESLD muscle tissue compared to healthy control muscle tissue using IPA, based on DEGs. Bars indicate the significance of the pathway activation/inhibition(left) and number of DEGs identified within the pathway (right). **C.** Top activated and **D**. Top inhibited cellular functions in ESLD muscle compared to healthy control muscle tissue, identified using IPA based on DEGs. **E.** Top activated upstream regulators in ESLD muscle compared to healthy control muscle tissue identified using IPA based on differentially expressed genes. n=18 Healthy control individuals. N=24 ESLD Patients. DEGs; Differentially expressed genes. ESLD; End stage liver disease. NS; Non-significant. FC; Fold Change. IPA; Ingenuity pathway analysis.

### ESLD aetiologies exhibit distinct skeletal muscle transcriptomic signatures

Next, considering ESLD can be subcategorised by aetiology (ArLD, MASLD and IALD), we investigated whether aetiology was associated with different skeletal muscle transcriptomic profiles (Figure 2A). We observed distinct skeletal muscle gene expression profiles for each of the disease aetiologies, with only 11 DEGs conserved across all aetiologies (Figure 2B). Some of these conserved DEGs (RABGGTB, IGFN1, PCGF5, DPYSL3) were consistent with those reported to display the greatest fold changes when considering ELSD as a collective cohort (Figure 1A). Subsequently, pathway analysis of DEGs for each aetiology indicated distinct differences in dysregulated canonical signalling pathways between groups (Figure 1C). For example, the MASLD transcriptome suggests metabolic dysfunction (oxidative phosphorylation, mitochondrial dysfunction) while ArLD may be associated with structural dysfunction within muscle (tight junction signalling, actin cytoskeleton signalling) and IALD more strongly linked with protein degradation (protein ubiquitination pathway, unfolded protein response). Similarly, upstream regulators predicted to drive the observed transcriptomic changes were also aetiology specific and included several pro-inflammatory cytokines including hepatocyte growth factor (HGF; MASLD subgroup) and interleukin-1 (IL-1;IALD and ArLD disease subgroups) (Figure 2D).

**Figure 2.**
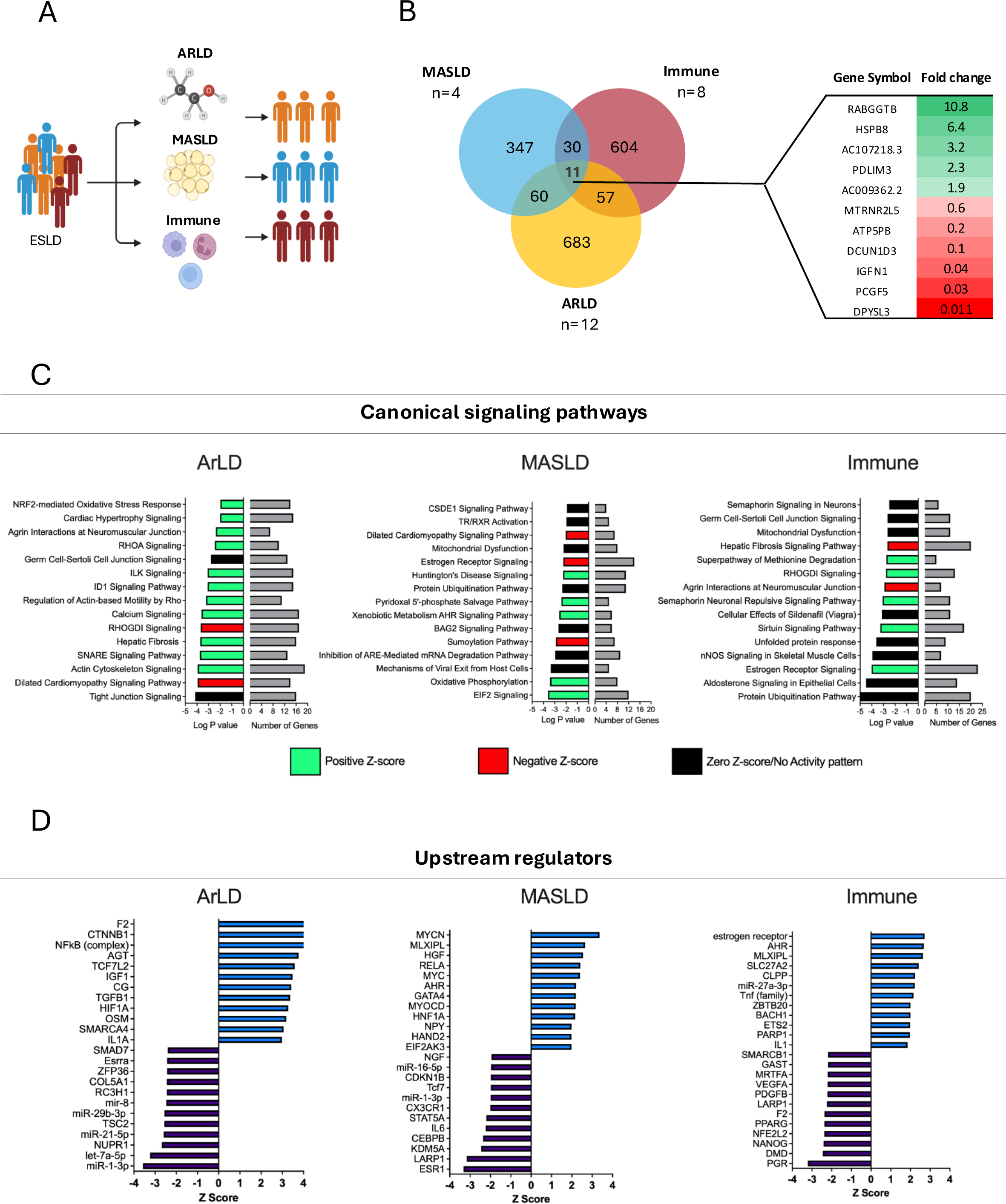
Skeletal muscle from ESLD patients exhibits a differential transcriptome depending on disease aetiology. **A.** Schematic overview of analysis, demonstrating how samples from the ESLD cohort were stratified for further analysis. **B.** Venn diagram highlighting the number of DEGs identified within the muscle tissue of ESLD patients sub-grouped by aetiology compared to appropriate age/sex matched control muscle tissue. DEGs common across all subgroups are indicated (right). **C.** Top canonical signalling pathways identified in the skeletal muscle for each ESLD aetiology compared to healthy control muscle tissue using IPA. Bars demonstrate the significance of the pathway and number of DEGs identified within the pathway. **D.** Top upstream regulators identified in the skeletal muscle for each ESLD aetiology compared to healthy control muscle tissue using IPA. ARLD; Alcohol Related Liver Disease. DEGs; Differentially expressed genes. ESLD; End-stage Liver Disease. MASLD; Metabolic Dysfunction-Associated Steatotic Liver Disease. Figure 2A was created in BioRender. Macleod, M. (2025) https://BioRender.com/k115sb3

### Patients with ESLD exhibit altered circulatory cytokines, with distinct aetiology-specific profiles that drive an atrophic phenotype in human muscle

In order to investigate whether circulating factors in ESLD may be driving the alterations observed in the skeletal muscle of ESLD patients, we transitioned to an *ex vivo* model in which primary human myotubes, obtained from a non-ESLD origin, were cultured in media containing 10% plasma from healthy controls, or individuals with ESLD for 24 h (Figure 3A). 24 h stimulation of primary human myotubes with ESLD plasma significantly (P=0.02) increased muscle protein degradation and significantly reduced (P=0.04) total protein in comparison to myotubes treated with healthy control plasma (Figure 3B). No significant effect on protein synthesis rate was observed (Figure 3B). Next, to determine the impact of soluble mediators on the muscle transcriptome, we performed RNA sequencing on primary human myotubes following 24 h treatment with control or ESLD patient plasma. Compared to myotubes treated with healthy plasma, ESLD plasma induced a significant transcriptomic shift (423 DEGs, P<0.05, Fold change±1.5) (Figure 3C). IPA analysis indicated that this resulted in modulation of canonical signalling pathways, similar to those identified in ESLD muscle tissue (Figure 1), including cachexia signalling and cellular senescence (Figure 3D). Upstream regulators predicted to drive these transcriptomic changes included increased pro-inflammatory cytokines (IL-1α) and a reduction in anti-inflammatory mediators (IL1-RA, IL-10) (Figure 3D). Upon dividing our dataset by disease aetiology, we again observed distinct myotube transcriptomic profiles, with only 68 common DEGs across all groups (Figure 3E). Plasma obtained from ArLD elicited the greatest impact on the myotube transcriptome (932 DEGs), followed by IALD disease (363 DEGs) and MASLD (136 DEGs). Similarly to the comparison between control and ESLD muscle tissue, DEGs induced upon ESLD plasma treatment of myotubes were associated with aetiology distinct dysregulation of canonical signalling pathways (ArLD: impaired cellular growth and division, MASLD: dysregulated metabolic pathways, IALD; inflammatory cytokines, neuronal impairment) and upstream regulators (Figure 3F). Additionally, overlap was observed when back translating these observations to corresponding muscle tissue pathway data (Figure 2).

**Figure 3.**
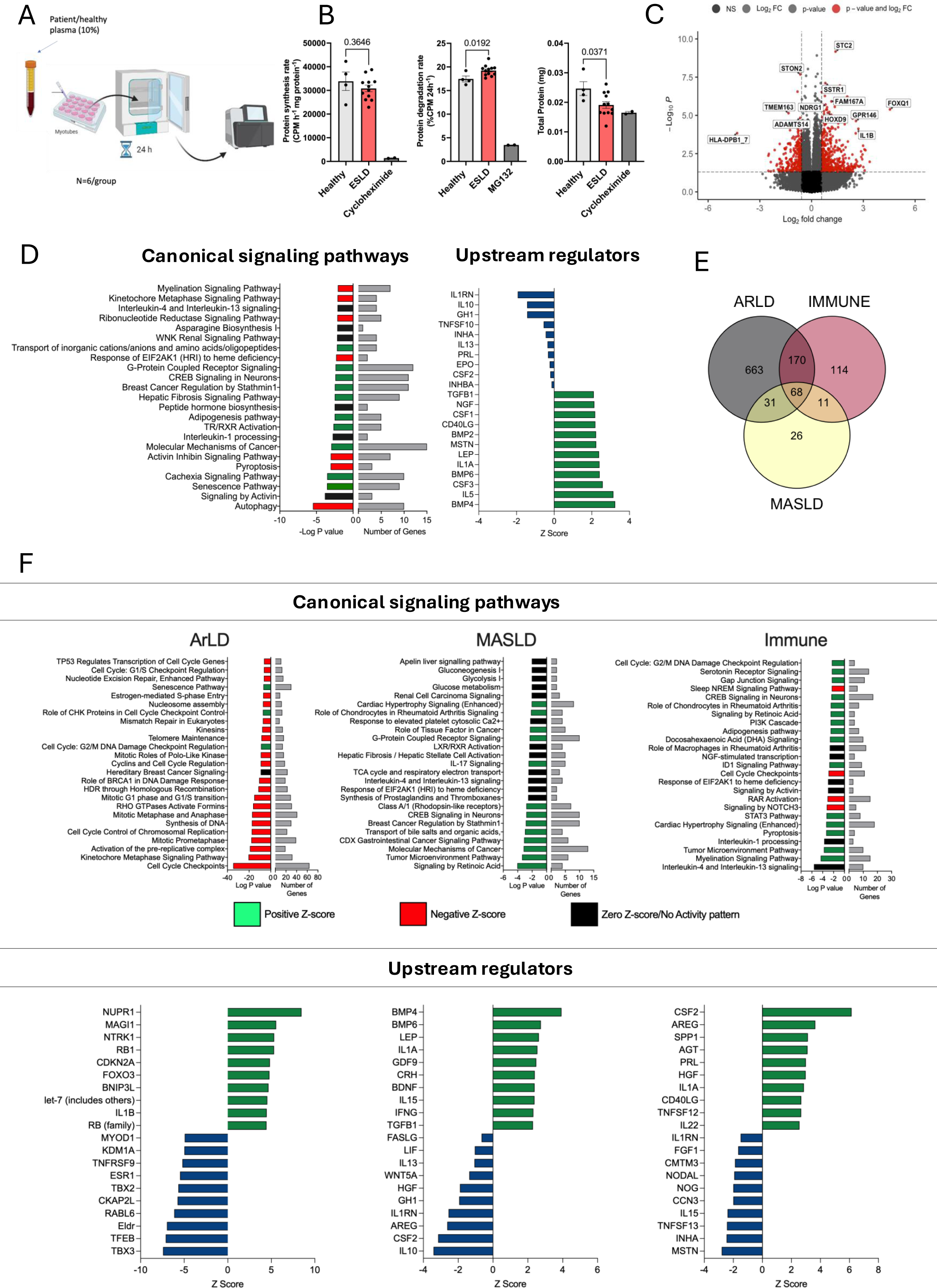
Plasma from ESLD patients drives aetiology dependent transcriptomic changes in human primary myotubes. **A.** Schematic overview of the experimental design used to generate data in figure 3. Primary human myotubes were treated with plasma from healthy individuals or ESLD patients for 24h, followed by RNA sequencing. **B.** Measurement of protein synthesis rate, degradation rate and total protein in human primary myotubes treated with ESLD patient plasma for 24h. **C.** Volcano plot of differentially expressed genes (P<0.05, Fold change >1.5) in primary human myotubes treated with ESLD patient plasma vs plasma from and age/sex matched healthy individuals. **D.** Left:Top canonical signalling pathways identified based on DEGs identified in primary human myotubes treated with ESLD patient plasma vs plasma from and age/sex matched healthy individuals for 24h, using IPA. Bars demonstrate the significance of the pathway and number of DEGs identified within the pathway. Right:Top upstream regulators (filtered on soluble mediators) based on DEGs identified in primary human myotubes treated with ESLD patient plasma vs plasma from and age/sex matched healthy individuals for 24h, using IPA. **E.** Venn diagram showing the number of DEGs identified within human primary myotubes following treatment with plasma from ESLD patients of different aetiologies vs appropriate age/sex matched healthy individuals for 24h. **F.** Top: Top canonical signalling pathways identified based on DEGs in primary human myotubes treated with ESLD patient plasma from different aetiologies vs plasma from appropriate age/sex matched healthy individuals for 24h, using IPA. Bars demonstrate the significance of the pathway and number of DEGs identified within the pathway. Bottom: Top upstream regulators identified based on DEGs in primary human myotubes treated with ESLD patient plasma from different aetiologies vs plasma from appropriate age/sex matched healthy individuals for 24h, using IPA For in vitro experiments, n=6 Healthy control plasma samples, n=6 ARLD Plasma samples, n=6 MASLD Plasma samples, n=6 IALD plasma samples. ARLD; Alcohol Related Liver Disease. DEGs; Differentially expressed genes. ESLD; End-stage Liver Disease. MASLD; Metabolic Dysfunction-Associated Steatotic Liver Disease. Figure 3A was created in BioRender. Macleod, M. (2025) https://BioRender.com/4548oyt

Having demonstrated a robust atrophic effect of ESLD patient plasma on primary human myotubes and identifying cytokines as potential mediators of this effect, we measured the concentration of 60 inflammation and age associated cytokines in the serum of patients with ESLD, compared to healthy controls. In total, 15 cytokines were found to be at a significantly higher concentration (P<0.05) and with 5 cytokines at a significantly lower concentration (P<0.05) in ESLD patient serum compared to healthy control serum (Table 1). PCA analysis indicated that healthy controls exhibited limited variation within their cytokine profiles, and such profiles were distinct from the majority of ESLD patients (Figure 4A). In-keeping with our transcriptomic data, IL1-RA was lower in ESLD patients (median, 85.2 pg/ml) compared to healthy controls (median, 113.0 pg/ml) (P=0.06). Additionally, levels of IL-1RA were positively associated with skeletal muscle mass (P=0.03, r=0.38) (Figure 4B) and negatively associated with liver frailty index (LFI), an integrative measure of physical function, (P=0.008, r=-0.42) (Figure 4C). Upon clustering of significant cytokines, we observed grouping of ESLD patients by aetiology (Figure 4D). Upon separating ESLD patients by disease aetiology, several cytokines were observed to be uniquely different (P<0.05) in comparison to healthy serum (Table 2), including greater IL-1α in the IALD subgroup (Figure 4E) (P=0.02), while HGF was significantly elevated in both MASLD and ARLD individuals (Figure 4E). Critically, this is in-keeping with aetiology specific upstream regulators inferred from our transcriptomic data (Figures 2-3). GDF-15 was *significantly* elevated (P<0.0001), across all disease aetiologies, with this increase being consistently 10-fold across all groups (Figure 4E).

**Figure 4.**
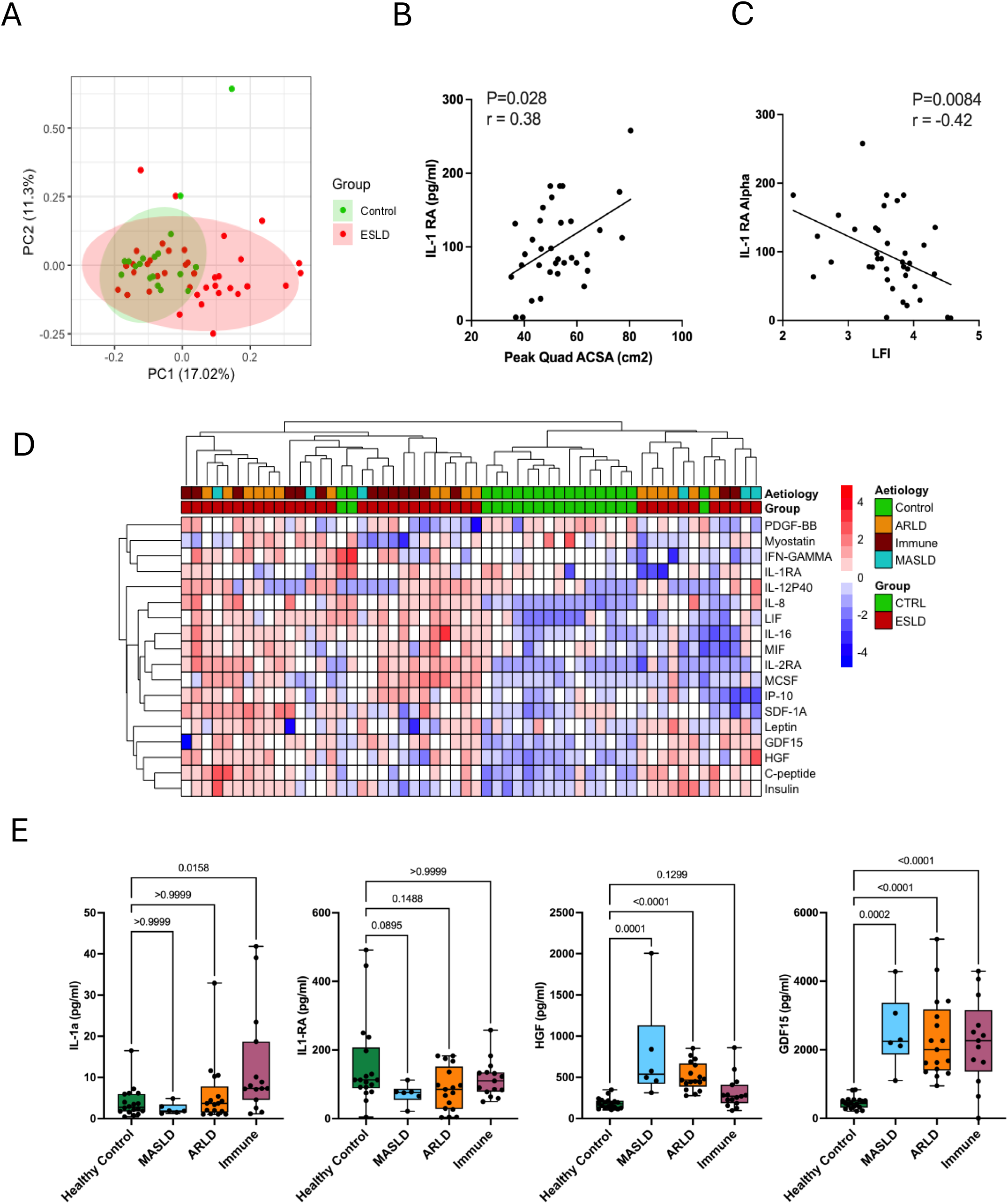
ESLD is associated with an altered circulating cytokine profile. **A.** PCA plot of all cytokines measured in the serum of ESLD patients (n=38) and healthy control individuals (n=18). **B.** Spearman correlation between peak quadricep ACSA and IL-RA in ESLD patients. **C.** Spearman correlation between peak quadricep ACSA and LFI. **D.** Heatmap displaying significantly different cytokines between ESLD patients and healthy control individuals. To generate the heatmap, raw cytokine values were expressed as a fold change and Log transformed. **E.** Concentrations of IL-1α, IL-1RA , HGF and GDF-15 in healthy control individuals and ELSD patients, separated by aetiology. ACSA, Anatomical cross-sectional area. ARLD; Alcohol Related Liver Disease. ESLD; End-stage Liver Disease. GDF-15; Growth Differentiation Factor15. HGF; Hepatocyte Growth Factor. IL-1RA; interleukin 1-Receptor antagonist. LFI, Liver frailty index. MASLD; Metabolic Dysfunction-Associated Steatotic Liver Disease.

**Table 1.** Significantly different cytokines in the serum of ESLD patients in comparison to healthy control individuals. Data are presented as median with lower and upper quartiles. statistical significance was determined by unpaired t-tests for parametric data and Mann-Whitney tests for non-parametric data. n=18 Healthy control individuals, n=38 ESLD unless otherwise stated. GDF-15, Growth/differentiation factor–15. HGF, Hepatocyte growth factor. IP-10, Interferon gamma-induced protein. LIF, Leukaemia inhibitory factor. MCSF, Macrophage colony-stimulating factor. PDGF, Platelet derived growth factor. PAI-1, Plasminogen activator inhibitor-1. SCF, Stem cell factor. SDF-1A, Stromal cell-derived factor 1.

**Table 2.** Significantly different cytokines in the serum of ESLD patients sub-grouped by disease aitiology compared to healthy control individuals. Data are presented as median with lower and upper quartiles. statistical significance was determined one-way ANOVA followed by Dunnett’s multiple comparison test for parametric data or Kruskal-Wallis test, followed by Dunn’s multiple comparison tests for non-parametric data. n=18 Healthy control individuals, n=6 MASLD, n=17 ArLD n=15 IALD unless otherwise stated. GDF-15, Growth/differentiation factor–15. HGF, Hepatocyte growth factor. IP-10, Interferon gamma-induced protein. LIF, (Leukemia inhibitory factor). MCSF, Macrophage colony-stimulating factor). PDGF, Platelet derived growth factor. PAI-1, (plasminogen activator inhibitor-1).

### Treatment of human myotubes with physiologically relevant doses of cytokines associated with ESLD promotes a sarcopenic phenotype

Serum GDF-15 was consistently elevated across all disease aetiologies and both HGF and IL-1/IL1-RA were implicated as upstream regulators in our muscle tissue and myotube RNA sequencing data and significantly different in ESLD patient serum. Subsequently we returned to our myotube model to investigate the role of these candidate cytokines on myotube growth, differentiation and metabolism, a key dysregulated pathway we identified in ESLD muscle (Figure 1). Regarding IL-1, we specifically, selected IL-1α due to its significant elevation in the plasma of the IALD cohort, while no significant differences in il-1β were identified across our cohorts.

Firstly, to investigate the impact of these candidate cytokines on myogenesis, we chronically stimulated human myoblasts during the course of their 8-day differentiation period (Figure 5A). Stimulation with a physiological dose of IL-1α (10 pg/ml) significantly reduced both myotube thickness (30%, P=0.02) and nuclei fusion index (P=0.03), and these effects were amplified following treatment with a higher dose (1 ng/ml) (Figure 5B-D). Similarly, GDF-15 significantly reduced myotube thickness (P=0.04), although it did not appear to impact upon nuclei fusion (Figure 5B-D). HGF did not impact either myotube thickness or nuclei fusion index following treatment with high or low doses (Figure 5B-D). Having observed an IL-1α-mediated effect on myogenesis, we next sought to confirm whether IL-1α impacted the expression of myogenic regulatory factors (MRFs), namely MyoD, myogenin (MYOG) and MYF5. Physiological doses of IL-1α significantly supressed the mRNA expression of both MYOG and MyoD at days 4 and 8 of myogenesis, in comparison to untreated controls (Figure 5E). IL-1α treatment had no effect upon MYF5 expression (Figure 5E). GDF-15 treatment significantly suppressed MYOG mRNA expression at day 4 (Figure S1). HGF also appeared to dysregulate myogenic gene expression, with significantly elevated MYOG expression observed at day 8 of differentiation with both physiological and supraphysiological doses (Figure S1).

**Figure 5.**
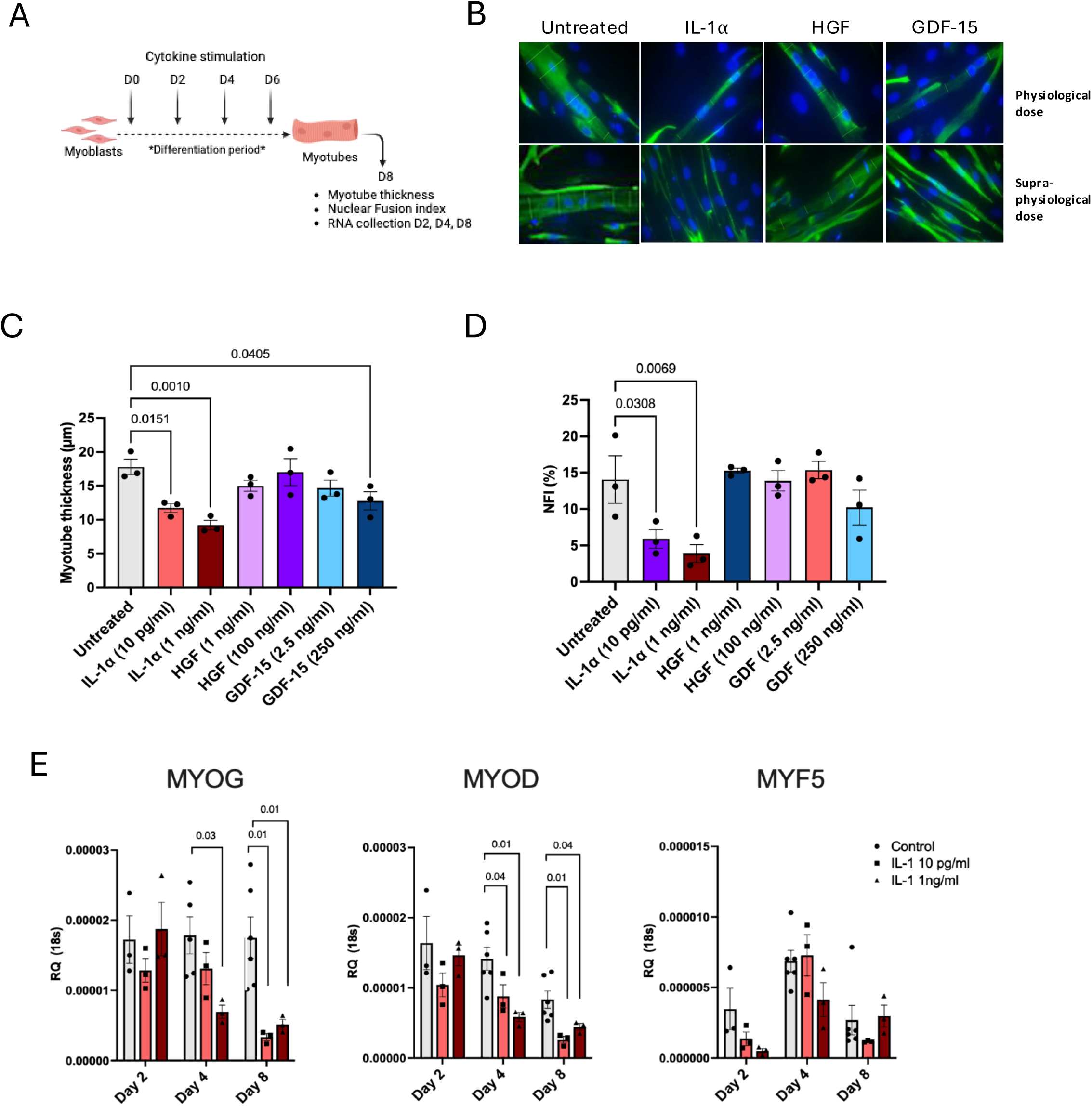
Physiological concentrations of candidate cytokines identified in the serum of ESLD patients impair differentiation of human primary myotubes. **A.** Schematic of experimental design. Confluent myoblasts were stimulated with recombinant IL-1 α, HGF or GDF-15 every 48 hours during routine media changes. Arrows indicate stimulation times and sample collection. **B.** Immunofluorescent staining of primary human myotubes at day 8 post differentiation following stimulation with or without recombinant human IL-1α, HGF or GDF-15. Desmin; green, DAPI; Blue. Images captured at 63x Magnification. **C.** Analysis of primary human myotubes thickness at day 8 post differentiation following repeated stimulation with or without recombinant human IL-1α, HGF or GDF-15. n=3 biological replicates per condition. **D.** Analysis of primary human myotubes nuclei fusion index at day 8 post differentiation following repeated stimulation with or without recombinant human IL-1α, HGF or GDF-15. n=3 biological replicates per condition. **E.** MYOG, MyoD and MYF5 mRNA expression in primary human myoblasts/myotubes following stimulation with or without recombinant human IL-1α for 2, 4 or 8 days. n=3 biological replicates per condition, except D4 and D8 untreated controls where n=6. Figure 5A was created in BioRender. Macleod, M.(2025). IL-1α Physiological dose=10pg/ml, supraphysiological dose=1ng/ml. HGF: Physiological dose=1ng/ml, supraphysiological dose=100ng/ml. GDF-15: Physiological dose=2.5 ng/ml, supraphysiological dose= 250ng/ml. https://BioRender.com/dmm4rzh, https://BioRender.com/ielva52.

We also examined the potential of these candidate cytokines to drive muscle atrophy by treating fully differentiated myotubes for 24 h (Figure 6A). Again, Recombinant IL-1α elicited the greatest atrophic effect, significantly reducing myotube thickness by 50% with a physiological dose (P=0.0003). Physiologically relevant doses of HGF and GDF-15 also significantly reduced myotube thickness (Figure 6B-C). In line with these findings, 24 h treatment of primary human myotubes with IL-1α upregulated the expression of atrophic genes (Fig 6D). IL-1α was also pro-inflammatory, with a robust and dose-dependent increase in IL-6 mRNA expression observed, which translated to greater secretion of IL-6 protein (Figure S2). A similar dose dependent upregulation of these atrophic genes was observed following treatment with recombinant HGF (Figure S3). No significant changes in myotube atrophic gene expression were observed following GDF-15 treatment (Figure S3).

**Figure 6.**
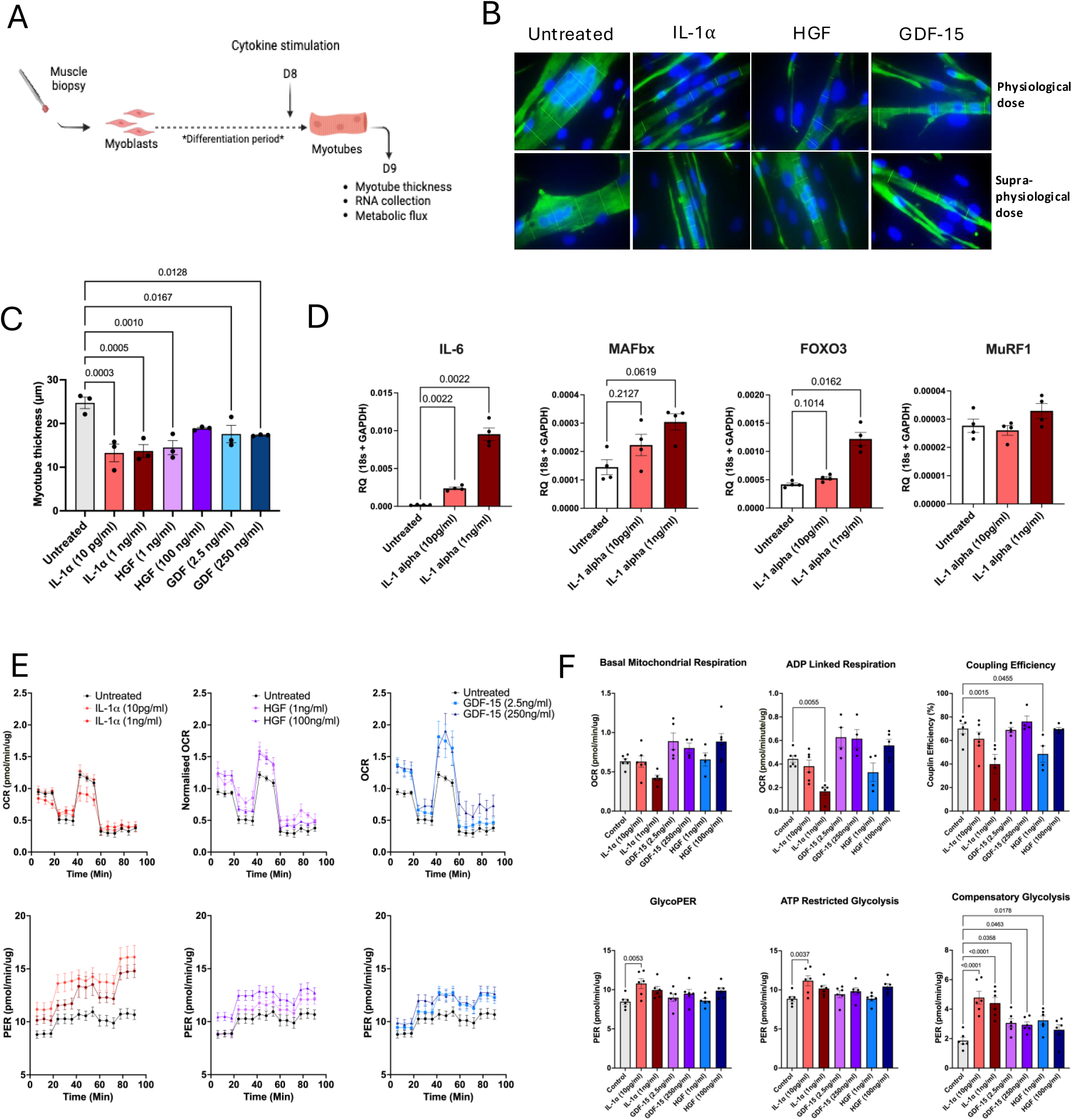
Physiological concentrations of candidate cytokines identified in the serum of ESLD patients cause atrophy of human primary myotubes. **A.** Schematic of experimental design. Differentiated myotubes were stimulated with recombinant IL-1α, HGF or GDF-for 24h at the end of the differentiation period. **B.** Immunofluorescent staining of primary human myotubes at day 8 post differentiation following stimulation with or without recombinant human IL-1 α, HGF or GDF-15 for 24 h at the end of the differentiation period. Desmin; green, DAPI; Blue. Images captured at 63x Magnification. **C.** Analysis of primary human myotubes thickness at day 8 post differentiation following stimulation with or without recombinant human IL-1α, HGF or GDF-15 for 24 h at the end of the differentiation period. n=3 biological replicates per condition. **D.** mRNA expression of atrophy associated genes; MAFbx, MuRF1 and FOXO3 and IL-6 in primary human myotubes following stimulation with or without recombinant human IL-1 for 24 h at the end of the differentiation period. n=4 biological replicates per condition. **E.** Top. OCR and Bottom. PER in primary human myotubes measured by Seahorse 96XF following stimulation with or without recombinant human IL-1α, HGF or GDF-15 for 24 h at the end of the differentiation period. OCR and PER measurements were normalised to total myotube protein. **F.** Analysis of seahorse metabolic flux data derived from OCR and PER measurements in response to oligomycin, BAM15, rotenone, antimycin A and monensin. n=6 per group, unless outliers were removed as indicated by individual data points. IL-1α Physiological dose=10pg/ml, supraphysiological dose=1ng/ml. HGF: Physiological dose=1ng/ml, supraphysiological dose=100ng/ml. GDF-15: Physiological dose=2.5 ng/ml, supraphysiological dose= 250ng/ml. OCR, oxygen Consumption rate. PER, proton efflux rate. Figure 6A was created in BioRender. Macleod, M.(2025).

Finally, to determine whether these candidate cytokines may also impact skeletal muscle function, we examined their impact on myotube metabolic flux, as mitochondrial dysfunction was highlighted to be dysregulated in our muscle and myotube transcriptomic data (Figure 1-3). 24 h treatment with physiologically relevant concentrations of IL-1α, HGF and GDF-15 were all identified to significantly perturb myotube metabolism (Figure 6E-F). Notably, IL-1α appeared to significantly impair mitochondrial oxidative phosphorylation and drive glycolysis in a dose dependent manner. In contrast, HGF and GDF-15 both appeared to significantly increase both oxidative phosphorylation and glycolysis, suggesting these cytokines also evoked metabolic stress (Figure 6E-F).

## Discussion

Here, we present the first evidence of significant transcriptomic changes in the skeletal muscle tissue of patients with ESLD. We demonstrate clear gene signatures associated with skeletal muscle atrophy and distinct mRNA profiles depending on the underlying disease aetiology of ESLD. Additionally, we highlight several candidate cytokines, with differential circulating levels in ELSD patients, and provide evidence that these cytokines drive transcriptomic and functional phenotypic changes in human myotubes consistent with the ESLD sarcopenic phenotype, including muscle metabolic dysfunction, muscle atrophy and impaired myogenesis.

ESLD profoundly impacted the human skeletal muscle transcriptome, which may underpin the skeletal muscle atrophy and impaired muscle function observed in these patients. Indeed, pathway analysis identified dysregulated mitochondrial function, inhibition of mTOR signalling, (a key pathway regulating skeletal muscle hypertrophy (20, 21) and increased activation of the protein ubiquitination system, (central to skeletal muscle atrophy (22) as being amongst the most differentially activated canonical signalling pathways in the skeletal muscle tissue from ESLD patients. Of note, the most significantly upregulated DEG within this dataset was DEPTOR, an inhibitor of mTOR (23). Sustained elevation of DEPTOR in the skeletal muscle of patients with ESLD may therefore contribute to the blunting of mTOR mediated protein synthesis and could be targeted to increase muscle anabolism (24). Conversely, top downregulated genes appear to be in keeping with protein catabolism and negative regulation of muscle mass. For example, IGFN-1 has recently been demonstrated to be necessary for fusion and differentiation of the murine myoblast cell line C2C12 (19, 25) and was similarly identified as a top downregulated gene associated with impaired muscular performance within aged skeletal muscle tissue (18). Downregulation of IGFN-1 with age is also in-keeping with increased cellular senescence being identified as a top cellular function within our dataset and our previous work demonstrating accelerated biological ageing of this liver disease patient cohort (26).

Upon sub-dividing our ESLD cohort based on disease aetiology, we identified a clear separation in the transcriptomic profile between groups, indicating that disease aetiology may differentially influence skeletal muscle gene expression. Additionally, we also report distinct transcriptional profiles in primary human myotubes following treatment with ESLD plasma from different aetiologies. Utilising murine models of MASLD and metabolic-dysfunction associated steatohepatitis (MASH), Guo et al similarly described distinct transcriptomic profiles within the muscle tissue of these two models of liver disease, with only 13.2% commonality amongst DEGs between MASLD and MASH groups (27). Such differential transcriptomic profiles may be explained by significant heterogeneity of circulating cytokines across aetiologies, as demonstrated in our data. Therefore, although ESLD is commonly associated with sarcopenia, targeted interventions based on disease aetiology may be critical to maximising their efficacy in improving muscle mass and function.

Here, we provide evidence for the cytokines IL-1α and HGF as central mediators of driving muscle atrophy in ESLD due to their identification as candidate upstream regulators of the ESLD muscle transcriptome, their atrophic effect on human muscle using an ex vivo myotube model, and their elevated concentrations in ESLD patient serum. IL-1 has previously been demonstrated to drive an atrophic phenotype, increase activation of NFKB signalling and impair fusion in C2C12 myotubes (28), whilst novel work by You et al. recently demonstrated that ablation of the NLRP3 inflammasome prevented skeletal muscle atrophy post denervation(29). There is more limited evidence for a role of HGF in skeletal muscle function, although under certain conditions, HGF may provide a protective role against muscle degeneration by stimulating satellite cell activation post-injury (30). Surprisingly, all of these data have been exclusively performed in animal models and rodent muscle cell lines and so there is very limited evidence of the impact of these cytokines on human muscle. To our knowledge, we present the first evidence of IL-1α and HGF in mediating atrophy in human primary myotubes, importantly at physiologically relevant concentrations. Therapeutically, there may be potential to repurpose existing drugs to target such cytokines, with anti-IL-1 therapeutics such as Canakinumab or Anakinra, which are already used for the treatment of rheumatoid arthritis and the auto-inflammatory disorders such as cryopyrin-associated periodic syndromes (CAPS) and TNF Receptor–Associated Periodic Syndrome (TRAPS). However, a recent pilot study adopting this approach reported no significant improvement of Anakinra on muscle protein metabolism in a relatively young cohort of haemodialysis patients (31). Our data also lends support for the development of aetiology specific interventions. For example, HGF was identified as an upstream regulator in MASLD skeletal muscle tissue and was also significantly elevated in the serum of these patients, but not the IALD subgroup. In contrast, IL-1 was predicted to drive transcriptomic changes in the IALDsubgroup, with IL-1α exclusively elevated in the serum of these patients.

Additionally, GDF-15 was consistently elevated across all aetiologies, suggesting an important biological role in ESLD patients. We recently reported an association of GDF-15 with accelerated biological ageing in ESLD muscle tissue (26) and others have recently demonstrated an association of GDF-15 with liver disease severity (32, 33). In addition, timely data from Groakre et al. demonstrated that ponsegromab, a novel monoclonal antibody targeting GDF-15, increased bodyweight, muscle mass and physical activity in a phase 2 double-blind, placebo-controlled study, supporting a role of GDF-15 in driving sarcopenia (34). Interestingly, the authors suggest beneficial effects may be mediated by central GDF-15 signalling in the brain, due to the localised nature of the GDF-15 receptor, GFRAL. Our data indicates GDF-15 may also directly act on muscle, and we have also observed similar expression levels of GFRAL in human myotubes as in neurones (Figure S4). This direct effect is supported by data from Zhang et al, who observed similar atrophic effects of GDF-15 in C2C12 myotubes, at similar dosages to those we used here (35). Collectively, these data indicate that therapeutics aimed at targeting circulating factors may present a novel treatment option to prevent skeletal muscle loss in ESLD in an aetiology specific manner.

### Limitations

A limitation of the present study was the small sample size and it will be important to confirm the present findings in a larger cohort and with studies specifically designed to examine aetiology-associated drivers of sarcopenia in ESLD. In addition, we have only interrogated circulating cytokines within the patient serum with the intention of identifying novel mediators of crosstalk influencing the skeletal muscle. However, the local interstitial skeletal muscle milieu may have a different composition to the circulation and so we may have also overlooked important cytokines associated with impacting the skeletal muscle. Finally, we only considered the role of cytokines in this study and so it will be important to elucidate the role of other circulating factors that may influence muscle function, including hormones, extracellular vesicles and metabolites.

## Conclusions

Patients with ESLD often present with sarcopenia, which is associated with reduced quality of life and increased mortality. Here, we present the first evidence of a dysregulated skeletal muscle transcriptome in ESLD patients, which was associated with atrophy and impaired muscle function. We also observed distinct muscle transcriptomic and differential serum cytokine profiles based on liver disease aetiology and demonstrated that candidate cytokines could drive an ESLD sarcopenia phenotype in human myotubes. Therefore, cohort-specific therapeutic interventional strategies may be the most effective approach to reduce sarcopenia in these patients.

## Acknowledgements

This study was funded by the National Institute for Health and Care Research (NIHR) Birmingham Biomedical Research Centre. The views expressed here are those of the authors and not necessarily those of the NIHR, NHS, or Department for Health and Social Care. Science Suite Inc. dba BioRender has granted Michael Macleod permission to use graphic in accordance with BioRender’s Terms of Service and Academic License Terms.

## Author Contributions

**Conceptualization**: TN, SLA, JML, CAG, MJA, LB, SWJ. **Formal analysis**: TN, SLA, MM, JP, MSS, KCM, JH. **Investigation**, TN, SLA, JIQ, AD, MM, JP, MSS, KCM, JH, CD, FRW. **Resources:** CAG, JML, LB, SWJ: **Data curation:** TN, SLA, SIQ, AD, MM, JP, JH, MSS, CD, KCM, FRQ. Writing –**original draft:** TAN, MM, MSS. **Writing, review & editing**: All. **Visualization:** TN, SLA, CAG, JML, LB, SWJ. **Supervision**: JML, LB, SWJ. **Funding acquisition:** TN, SLA, AME, MJA, CAG, JML, LB, SWJ.

## Competing interests

The authors declare no competing interests

---

**Supplementary Figure 1.**
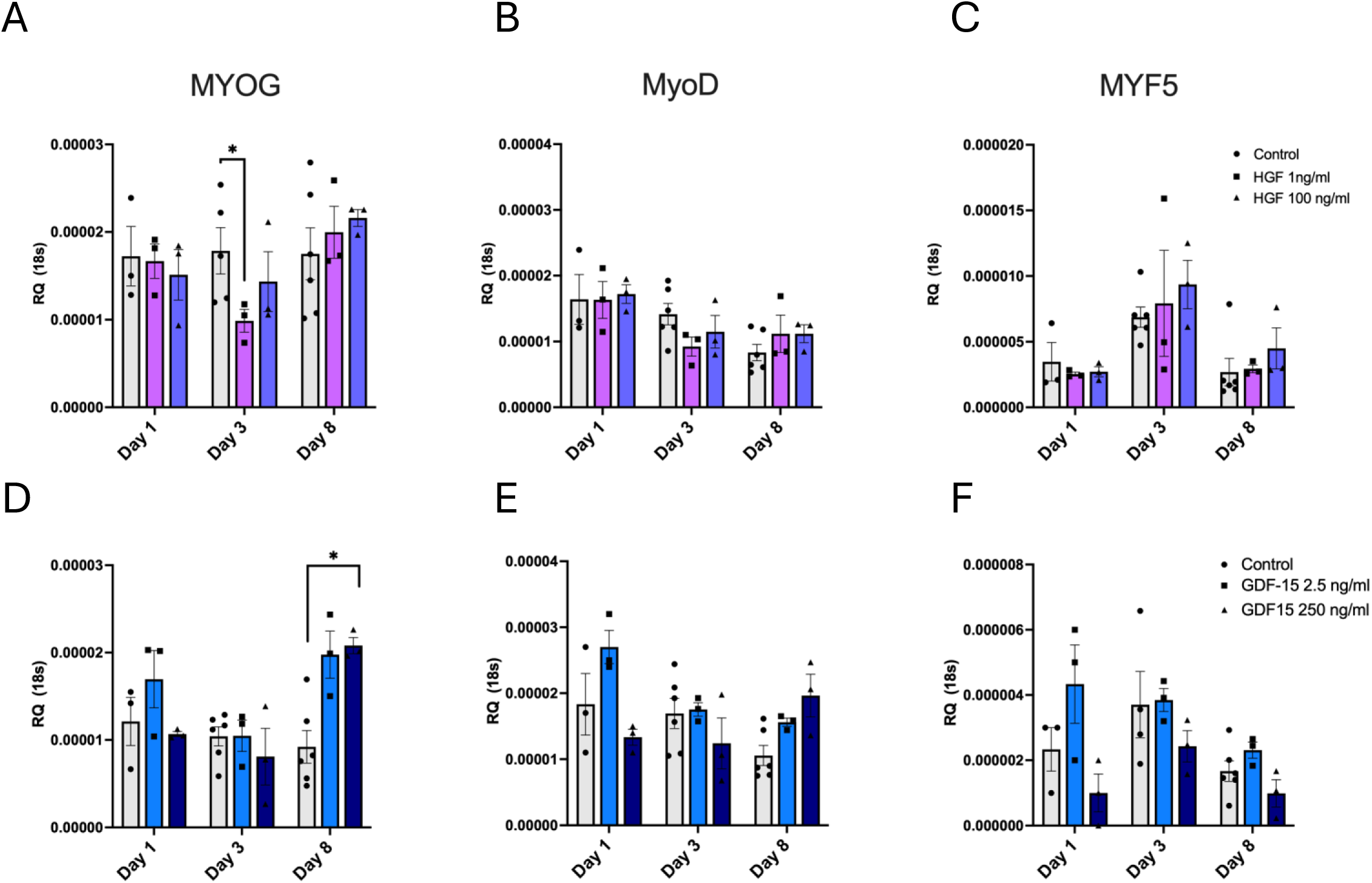
MYOG, MyoD and MYF5 mRNA expression in primary human myoblasts/ myotubes determined by qRT-PCR, following stimulation with or without recombinant human HGF (A-C), or GDF-15 (D-F) for 2, 4 or 8 days. n=3 biological replicates per condition except D4 and D8 untreated controls where n=6. * Denotes P value < 0.05).

**Supplementary Figure 2.**
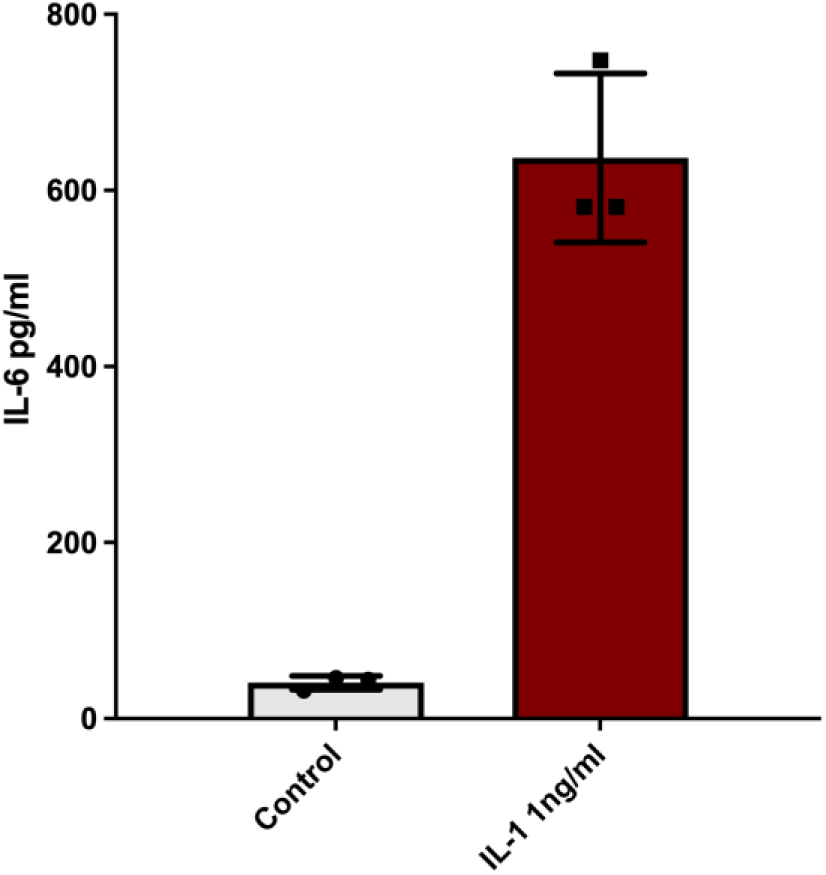
Secretion of IL-6 protein, from primary human myotubes following 4h stimulation with IL-1, measured by ELISA. Culture media was switched to serum free media upon starting cytokine stimulations, to avoid measurement of IL-6 derived from FBS. N=3 patient replicates.

**Supplementary Figure 3.**
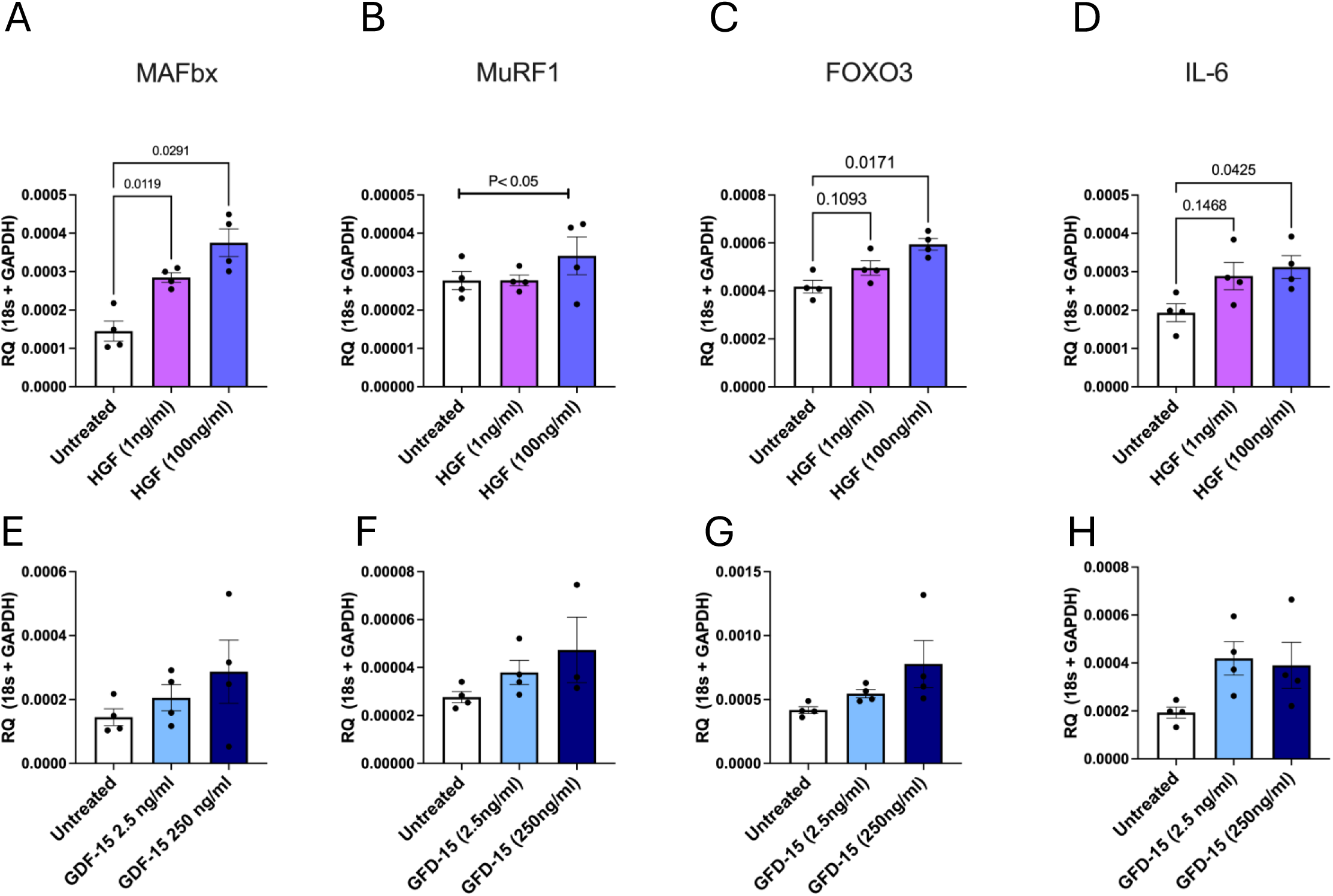
mRNA expression of atrophy associated genes; MAFbx, MuRF1 and FOXO3 and IL-6 in primary human myotubes following stimulation with or without recombinant human HGF (A-D) or GDF-15 (E-H) for 24 h at the end of the differentiation period. n=4 biological replicates per condition.

**Supplementary Figure 4.**
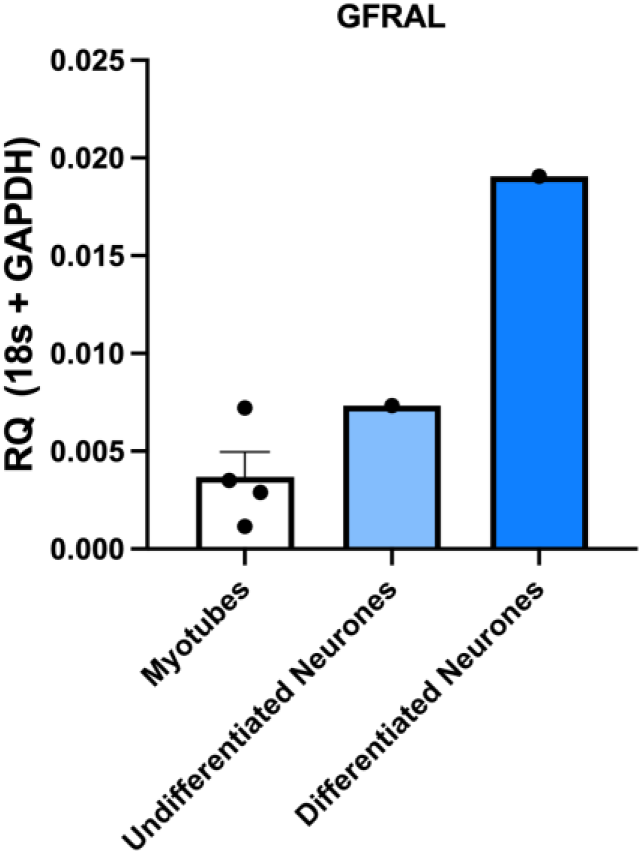
GFRAL mRNA expression in primary human myotubes (n=4 patient replicates) and undifferentiated and differentiated neurones.

## Notes

This work was supported by funding from the NIHR Birmingham Biomedical Research Centre, Birmingham, UK

### Competing Interest Statement

The authors have declared no competing interest.

